# LSD1 inhibition improves efficacy of adoptive T cell therapy by enhancing CD8^+^ T cell responsiveness

**DOI:** 10.1101/2023.10.27.564395

**Authors:** Isabella Pallavicini, Teresa Maria Frasconi, Elena Ceccaci, Silvia Tiberti, Carina B Nava Lauson, Elisa Preto, Alberto Bigogno, Marta Mangione, Eleonora Sala, Matteo Iannacone, Mirela Kuka, Luigi Nezi, Saverio Minucci, Teresa Manzo

## Abstract

The lysine-specific histone demethylase 1A (LSD1) has been described to play a role in antitumor immunity; however, the role of LSD1 in shaping CD8+ T cell (CTL) differentiation and function is not understood. Here, we showed that pharmacological inhibition of LSD1 (LSD1i) in CTL elicited phenotypic and functional alterations of the CTL that led to robust antitumor immunity in the context of adoptive T cell therapy (ACT). In addition, the combination of anti-PDL1 therapy with LSD1i-based ACT resulted in a complete tumor eradication and long-lasting tumor-free survival. This study demonstrated that LSD1i together with anti-PDL1 therapy complement each other’s deficiencies and produce a better tumor response in a melanoma model, in which both immune and epigenetic therapy alone have shown limited efficacy. Collectively, these results set the translational potential of modulating LSD1 to improve antitumoral responses generated by ACT and anti-PDL1 therapy, which provide a strong rationale for a combination trial of LSD1i and immunotherapy.

## INTRODUCTION

Adoptive cell therapy (ACT) harnesses the full potential of CD8+ T cells (CTL) to recognize tumor cells and carry out an anti-tumor effector function. This personalized therapy has shown promising results on different tumor types (Kalos et al., 2011; Brentjens et al., 2013; Maude et al., 2014; Rapoport et al., 2015) and clinical trials are being conducted worldwide to further implement it. However, the response of solid tumors to ACT is still marginal (Young et al., 2022)

ACT clinical success is currently hampered by: the acquisition of a dysfunctional state during the *in vitro* expansion step of T cells, which limits their survival and persistence after infusion into the patient (McLane et al., 2019); the loss of metabolic functional plasticity of T cells, which is fundamental to adapt and survive in the hostile TME (Kishton et al., 2017; Manzo et al., 2020; Nava Lauson et al., 2023) and the immunosuppressive influence of the tumor microenvironment (TME) (Salas-Benito et al., 2023). This, longevity, persistence and functionality are critical determinants for the efficacy of T cell-based immunotherapies (Klebanoff et al., 2012; Sukumar et al., 2017), implying that the phenotype of T cell products can profoundly impact therapy efficacy (Mondino and Manzo, 2020).

In this scenario, maintaining T cells in a less-differentiated state and with high plasticity and fitness during *ex vivo* T cell production and after infusion may have a strong therapeutic impact. Recent findings suggest a major role for epigenetic programs in determining and maintaining T cell fate decisions (Gray et al., 2017; Goswami et al. 2018). T cells may resist current strategies for reversing T cell exhaustion as a result of stable changes in gene regulation due to epigenetic reprogramming (Philip et al., 2017; Sen et al., 2016; Pauken et al., 2016). For instance, progressive de novo DNA methylation is an essential process inaugurating CD8+ T cell exhaustion. This results in a distinct gene repression signature that is preserved during PD-1 blockade (Ghoneim et al., 2017). Moreover, disruption of the *TET2* gene (encoding an enzyme that regulates DNA demethylation) in CAR-T cell resulted in clonal expansion of a single CAR-T cell that induced leukemia remission (Fraietta et al., 2018). Thus, manipulating the activity of demethylating enzymes that contribute in establishing exhaustion and memory programs may hold a key to obtaining a long-lived pool of engineered T cells with sustained anti-tumor responses, implementing the use of ACT in oncology.

LSD1 is a lysine-specific histone demethylase 1A highly expressed in several cancers - such as prostate, breast and hepatocellular-where it is associated with poor prognosis (Hosseini and Minucci, 2017). Interestingly, recent studies have underscored a role for LSD1i in influencing anti-tumor immunity (Sheng et al., 2018; Qin et al., 2019) by promoting an epigenetic program that modulate exhausted CTL (Liu et al., 2021; Tu et al., 2020). However, the impact of perturbing T cell-intrinsic LSD1 on CTL differentiation and function as well as their anti-tumoral responses have not been fully explored.

Here, we provided evidences that *in vitro* epigenetic reprogramming of CTL with LSD1i imprints a less-differentiated fate and ameliorates T cell exhaustion as manifested by inhibitory receptor reduction and enhanced metabolic fitness. As a direct consequence, LSD1-instructed T cells showed enhanced persistence and antitumor effects in a murine melanoma model of ACT. Furthermore, a rationally-designed combination of anti-PDL1 and LSD1i therapy resulted in a complete tumor eradication and long-lasting tumor-free survival. In conclusion, our study demonstrates that LSD1 could be considered as a new actionable target to fine-tune CTL functionality and fitness maintenance, linking CTL epigenetic tweaking to anti-tumor surveillance. This pave the way for a new generation of immunotherapies, where LSD1 inhibition can be used to potentiate the immunotherapy efficacy.

## RESULTS and DISCUSSION

### LSD1i *ex vivo* promotes a memory-like phenotype in CTL

LSD1i supports the anti-tumor immune response to anti-PD1 therapy by enhancing effector functions of tumor infiltrating CTL (Sheng et al., 2018; Qin et al., 2019; Liu et al., 2021). However, the role of LSD1i in regulating normal T cell activation and differentiation is yet to be determined. To understand the mechanism by which LSD1 influences CTL responsiveness, we examined the direct effect of LSD1 on CTL fate and function by activating CTL in the presence of LSD1 inhibitor (MC_2580 1) (Binda et al., 2010) at a final concentration of 2μM and compared them with controls (LSD1i-versus CTR-CTL) (experimental scheme, Figure 1A). Having found that LSD1i did not compromise overall viability of CTL (Figure S1A), we evaluated its effect on the proliferation and activation of CTL. LSD1i-CTL and CTR-CTL hold the same rate of proliferation - as assessed by the CFSE *in vitro* assay (Figure S1B) - and the same extent of activation – as demonstrated by similar cell surface marker expression (Figure S1C).

**Figure 1.**
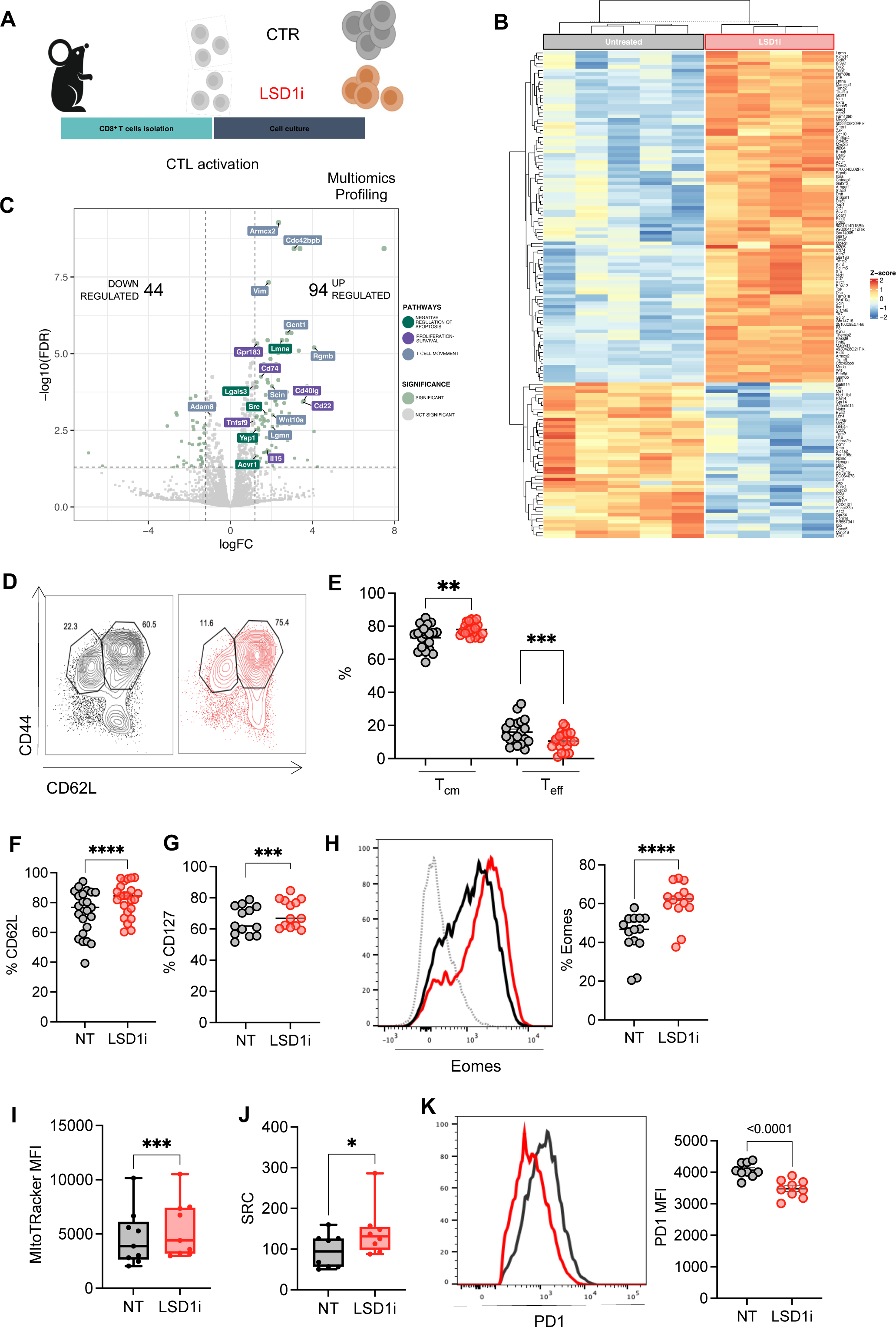
LSD1i treatment promotes a memory-like phenotype in CTL. **A.** Experimental design. CTL were isolated from spleen and lymph nodes of C57B6 mice and activated with anti-CD28, anti-CD3 and IL2 in presence or absence of LSD1i for 72h**. B.** Heatmap showing DEGs in LSD1i and CTR-CTL as assessed by RNA-seq (FDR< 0.05 and linear logFC > 1.2). Data from 4 to 5 mice for each group are shown. **C.** Volcano plot showing DEGs in LSD1i and CTR-CTL. Dot size reflects the significance of each genes. DEGs (FDR< 0.05 and linear logFC > 1.2) are highlighted in light green. Label color code identify genes from different pathways found significatively enriched in IPA analysis. **D-H.** Memory characterization measured as percentage of memory CD44^high^ CD62L^high^ (T_CM_) and effector CD44^high^ CD62L^low^ (T_EFF_) cells (D-E, n=24); expression of CD62L (F, n=24), CD127 (G, n=14) and Eomes (H, n=14). **I-J.** Metabolic profile assessed by MitoTracker (I, n=8) and Spare Respiratory Capacity (SRC, J, n=8). **K.** PD1 expression levels (n=9)

To better understand the impact of LSD1i treatment on CTL differentiation, we performed a transcriptional analysis of LSD1i-CTL, which highlighted 138 (44 down-regulated and 94 up-regulated) differentially expressed genes (DEGs) (Figure 1B-C; Table S1). Among the upregulated ones, we detected genes related to CTL memory differentiation pathways (Figure S1D), such as *Sell* and *Il15*, which mark long-lived memory cells (Martin and Badovinac, 2018), the transcription factor *Eomes* - key player in memory homeostasis (Doering et al., 2012), *Ccr10* and *Klrc1*-associated with optimal formation of memory T cell populations (Zaid et al., 2017). In line with the induction of a memory signature, LSD1i also modulated genes that are associated with a positive regulation of CTL proliferation and survival (i.e. *Gpr183*, *Tnfsf9*, *Cd40lg*, *Cd22, Il15*; Figure 1C) as well as negative regulation of apoptosis (i.e. *Src*, *Lgals3*, *Acvr1*, *Yap1*; Figure 1C), suggesting that LSD1i promotes survival of CTL. Of note, RNAseq analysis also showed a significant upregulation of genes involved in T cell migration and cytoskeleton remodeling, suggesting a possible advantage for LSD1-CTL to better be recruited to the tumor site (i.e. *Armcx2*, *Cdc42bpb*, *Lamna*, *Rgmb*, Wnt10, Adam10; Figure 1C). These data pointed out a role for LSD1i in skewing CTL toward a memory-like phenotype, a finding that was independently validated by employing multiparametric flow-cytometry. Indeed, LSD1i-CTL had reduced frequencies of short-lived terminal effector CTL - identified as CD44^high^CD62L^neg^ - paralleled by increased frequencies of long-lived central memory cells, identified by CD44^high^ CD62L^high^ (Figure 1D-E). Likewise, LSD1i favors the expression of canonical memory-associated markers - like CD62L (Figure 1F) and CD127 (Figure 1G) - increases the protein levels of the transcription factors Eomes (Figure 1H), which promotes a memory differentiation (Doering et al., 2012), and lowers the expression of the effector-associated transcription factor Tbet (Figure S1E). Thus, we can conclude that LSD1i promotes a memory-like phenotype in CTL. Furthermore, since memory T cell hold a superior mitochondrial spare respiratory capacity (SRC) allowing for improved functionality under stressed conditions (van der Windt et al., 2012), we determined whether LSD1i impacts on CTL cellular metabolism. We observed a significant increase in mitochondrial membrane potential upon LSD1i treatment, as assessed by the increased quantification of MitoTracker Orange staining (Figure 1I). Consistently, analysis of their bioenergetic profile highlighted that LSD1i-CTL harbor a significant higher SRC (Figure 1J), potentially allowing them to better cope with stressful conditions. Overall, these data indicate that LSD1i treatment confers to CTL an improved metabolic fitness, typical of memory T cells. Strikingly, despite showing similar activation status (Figures S1C), LSD1i-CTL displayed a significant drop in PD1 expression (Figures 1K), which might counteract the development of an exhausted phenotype within the TME. Furthermore, we confirmed that LSD1i treatment was able to shape towards a less differentiated memory phenotype also antigen-specific T cells. To this end, we used CTL from OT1 and P14 mice (expressing a transgenic TCR that recognizes OVA and LCMV GP antigens, respectively) and activated them using the cognate peptide in the presence of absence of LSD1i. LSD1i proved capable of inducing Eomes upregulation and PD1 downregulation also in antigen-specific CTL (Figure S1F-I). Thus, we concluded that LSD1i might redirect CTL away from exhaustion and toward a memory-like phenotype with increased metabolic fitness, potentially improving the likelihood to survive within a harsh TME.

### *Ex vivo* LSD1i enhances efficacy of ACT therapy

To evaluate whether the features conferred to CTL *in vitro* by LSD1i enable them to better adapt to the inhibitory TME, persist longer and mediate a durable tumor control, we employed an ACT model for established B16-OVA melanoma tumors by generating OVA-specific T cells *ex vivo* in the presence or absence of LSD1i (LSD1-ACT and ACT, respectively) and injecting them intravenously in tumor-bearing mice (experimental scheme in Figure 2A).

**Figure 2.**
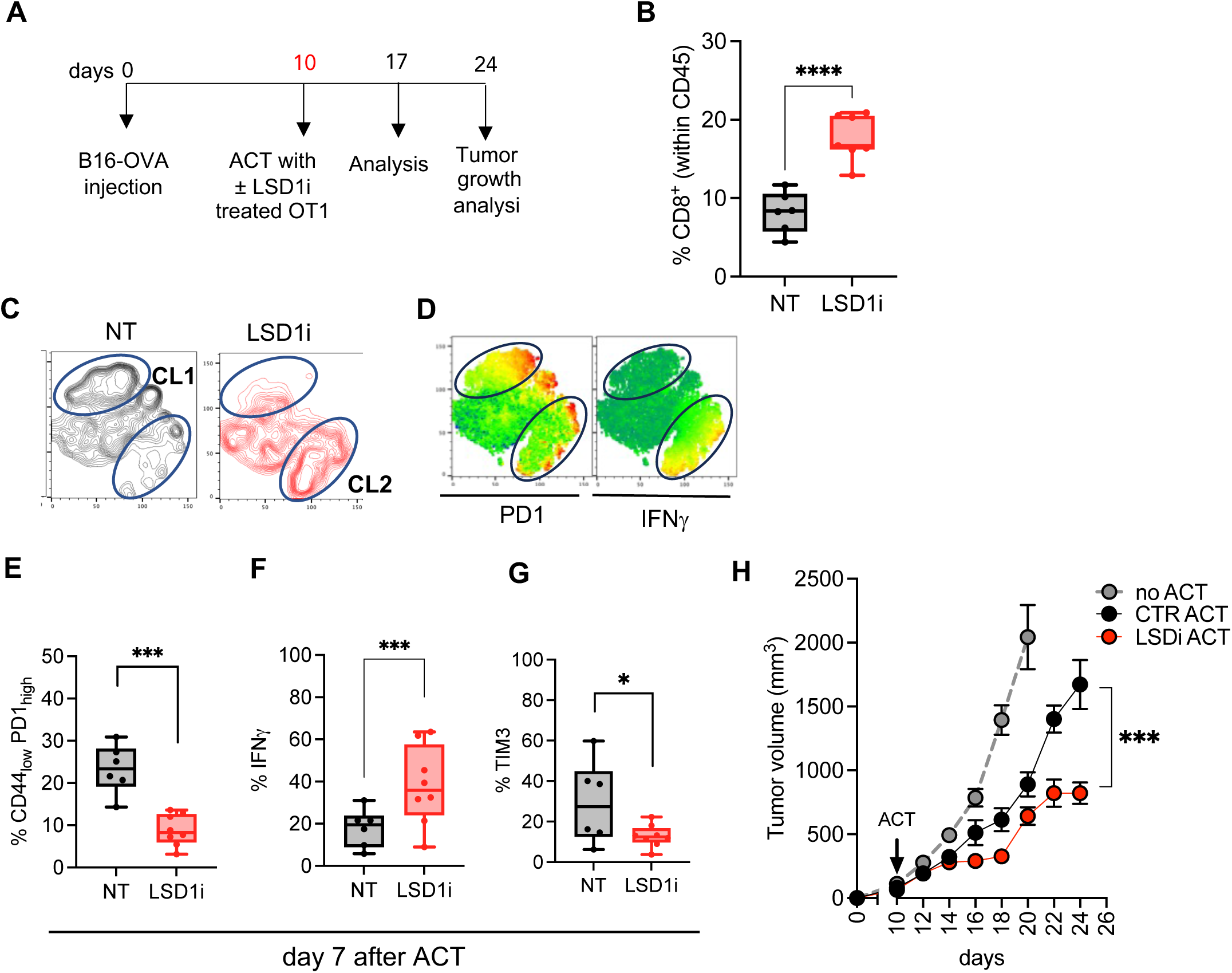
LSD1 inhibition during *ex vivo* T cell expansion increases ACT efficacy. **A.** ACT experimental design. **B-G.** Analysis of the CTL infiltrate 7 days post-infusion. Frequencies of CD8^+^ (within total CD45^+^) infiltrating the tumor (B), tSNE representation based on PD1 and IFNγ (C-D), exhaustion profile assessed as PD1_high_CD44_low_ (E), frequencies of IFNγ secreting CTL (F) and TIM3^+^ CTL (G). **H.** Tumor growth curve (n = 23 for CTR-ACT, 21 for LSD1iACT, and 11 for No-ACT; 4 independent experiments).

The analysis of the tumor infiltrating immune cells - one week after ACT - showed a significant enrichment of CTL in tumor from mice treated with LSD1i-CTL (Figure 2B), luckily due to their improved ability to migrate (Figure 1B) and to survive better because of their memory phenotype (Figure 1D). High-dimensional flow cytometry enables a comprehensive assessment of the phenotypic diversity between LSD1- and CTRL-CTL. t-distributed stochastic neighbour embedding (tSNE) analysis confirmed that CTL profiles were substantially different in the presence or absence of LSD1i (Figure 2C) and identified two CTL clusters (CL): CL1, that is characterized by high PD1 and low IFNγ levels and overrepresented in CTRL-CTL, and CL2 instead characterized by lower PD1 expression and higher IFNγ expression and specifically enriched in LSD1i-CTL (Figure 2D). Manual gating strategy revealed that, compared to tumor from mice that received a control ACT, those from mice transferred with a LSD1i-ACT featured CTL with a less-exhausted phenotype, as indicated by the lower frequencies of CD44^low^/PD1^high^ CTL (Figure 2E), higher ability to secrete IFNg (Figure 2F) as well as decreased TIM3 expression (Figure 2G). This analysis indicates that the advantage of a less-differentiated phenotype given *in vitro* by LSD1i prior to transfer favors a quick and strong recall *in vivo* 7 days after transfer. Finally, all these features enable LSD1-ACT to control tumors more efficiently when infused in tumor-bearing mice (Figure 2H and S2A-B), demonstrating their superior therapeutic potential. Of note, CTR-CTL were able to significantly control tumor growth, in line with the high therapeutic efficacy of ACT approaches.

However, LSD1i was able to further improve the anti-tumor potential of CTL. Thus, we conclude that LSD1i enhances the ACT therapy efficacy. Thus, we underscore a new role for LSD1i in CTL anti-tumor response being able to modulate CTL differentiation towards a more functional phenotype that operates a more potent acute tumor debulking.

### LSD1i enforces an IFN-γ/PD-L1 axis between T cells and tumor

We next wondered whether the LSD1i treatment induced a stable and persistent functional rewiring in CTL. Analysis of the tumor infiltrating CTL at the time of sacrifice, did not show any substantial advantage of the LSD1i-ACT groups. Indeed, frequencies of IFNγ-secreting CTL were comparable between the two groups (Figure S2C) and, surprisingly, PD1^+^ “exhausted” CTL were even significantly more abundant in LSD1i-CTL, when compared to CTR-CTL (Figure S2D-E). Thus, despite the significant improvement in the acute tumor control, LSD1i-induced reprogramming in CTL did not persist overtime.

We next asked about the potential mechanism whereby LSD1i-CTL would reacquire a dysfunctional phenotype typical of tumor-infiltrating CTL. When performing an IPA upstream regulator analysis to identify upstream regulator and predict whether they are activated or inhibited based on the observed gene expression changes in the RNAseq dataset, we identified *IFN*γ as the most significative activated regulator, as well as its upstream regulator *Sox4* and *IL12* (Figure 3A), suggesting an emergency of *IFN*γ signature in the iLSD1-CTL (Kuwahara et al., 2012; Trinchieri, 2003).

**Figure 3.**
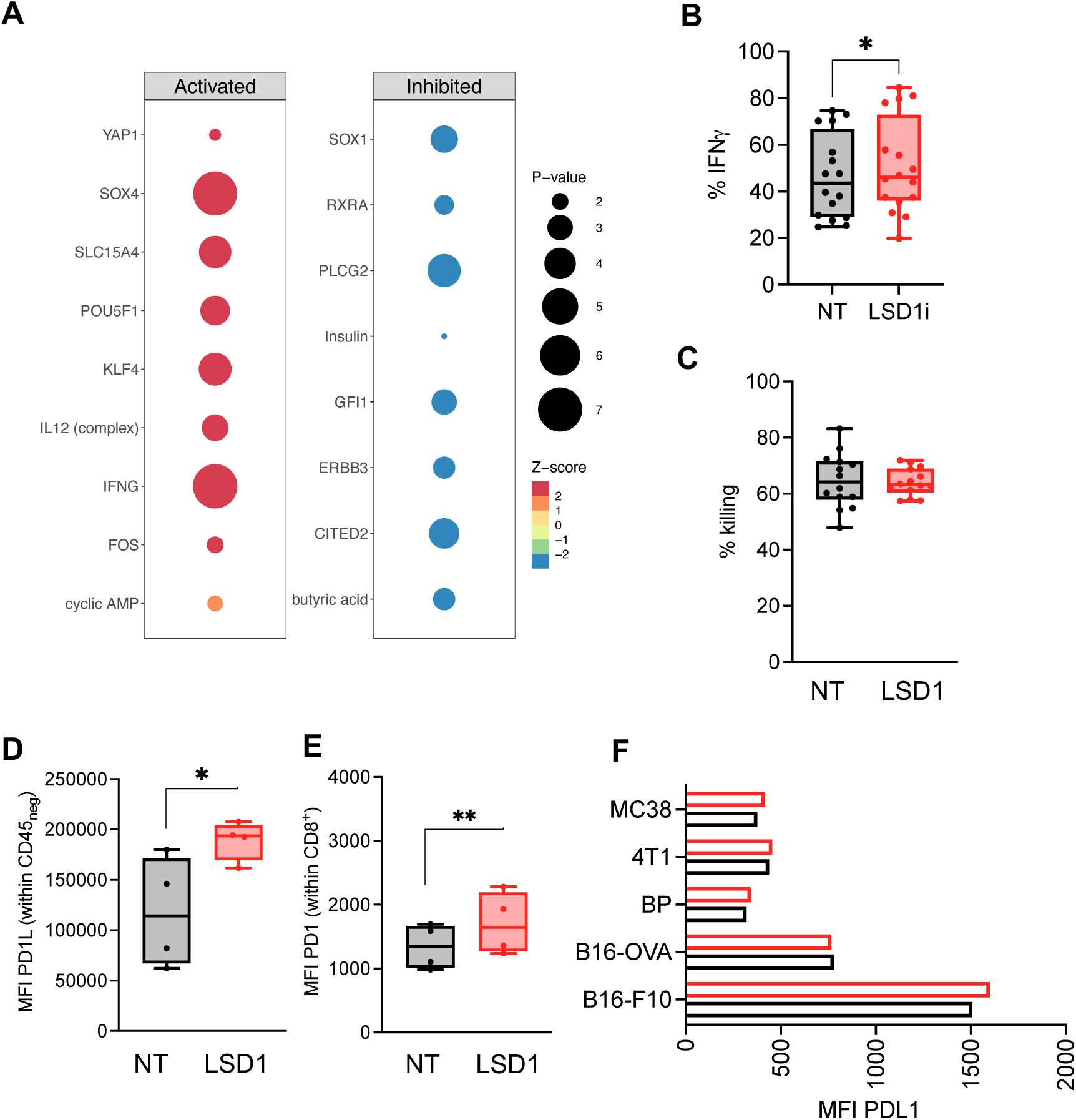
LSD1i treatment induces up-regulation of PD1/PDL1 axes. **A.** Bubble plot showing IPA prediction of upstream regulator TF based on DEG resulted from RNAseq analysis. TF are divided based on the prediction of Activated or inhibited: size represent the - log10(Pvalue) while color indicate the Zscore. **B.** IFNγ production and **C.** killing capacity of CTL±LSD1i (n=11). **D-E.** PDL1 (D) and PD1 (E) expression on B16-OVA tumor cells and CTL, respectively, in co-culture 1:1 (n=4). **F.** PDL1 expression on B16-OVA tumor cells ±LSD1i (n=4).

This result was independently confirmed at protein levels - as demonstrated by the higher amount of IFNγ produced by LSD1i-CTL (Figure 3B). To assess whether higher levels of IFNγ had functional consequences, we probed the antigen-specific antitumor killing capacities of LSD1i– or CTR-CTL in an *in vitro* killing assay (Figure 3C), finding that LSD1i- and CTR-CTL exhibited the same ability to kill cancer cells.

Higher IFNγ levels have been shown to have an opposite effect on a TME. If from one side it is needed to sensitize the tumor cells to be killed (Grasso et al., 2020; Hoekstra et al., 2020; Thibaut et al., 2020), from the other side it can induce PDL1 up-regulation in the TME, which promotes PD1 up-regulation in CTL and decreases their effector function (Benci et al., 2016; Paschen et al., 2022). To test this hypothesis, we measured PDL1 expression in tumors that were exposed to LSD1i- or CTR-CTL. Interestingly, we observed that tumors co-cultured with LSD1i-CTL expressed significant higher levels of PDL1 (Figure 3E), which was paralleled by PD1 up-regulation in CTL (Figure 3F). Of note, this was specifically induced by ILSD1i-treated CTL because LSD1i alone on tumor cells do not induce PDL1 upregulation (Figure 3G). Thus, we can conclude that LSD1i enforces an IFN-γ/PD-L1 axis between T cells and tumor, which explains the loss of the functional advantages given *in vitro* by LSD1i.

### The combination of LSD1i-based ACT and anti-PDL1 therapy promotes long-lasting anti-tumor response and establish immunosurveillance

Our data suggested that LSD1i-CTL initiates both a negative and a positive feedback loop. It elicits an IFNγ response in tumors that promotes tumor attack from CTL but also induces PD-L1 expression on tumor cells that blunts CTL activity by binding to its receptor PD1, thereby recreating an immune-suppressive TME, which in turn favors the development of an exhaustion state in CTL. Therefore, we speculated that inhibition of PD-L1 may be key to prevent the exhaustion of CTL promoted by a counterregulatory mechanism involving enhanced IFNγ production in the TME. To this aim, we decided to combine ACT regimen with anti-PDL1 and evaluate whether the combination increases effectiveness of ACT compared to monotherapies. Although the addition of an anti-PD-1 to a CTR-ACT provided only minimal benefits on top of CTR-ACT in the B16-OVA melanoma model (Figure S3A), we found that the combination of LSD1i-ACT and anti-PDL1 therapy drastically reduced tumor growth (Figure 4A-B and S3A) and, most importantly, it increased long-term survival compared with anti–PDL1 or ACT monotherapy (Figure 4C). Analyzing the tumor-infiltrating CTL, we noted that the combination therapy with LSD1i-ACT and anti-PDL1, as compared with monotherapy with LSD1i-ACT or anti-PDL1, reduced the percentage of exhausted CTL expressing PD1 or TIM3 inhibitory receptor (Figure S3B-C) and resulted in an increased infiltration of tumor-specific CTL (Figure S3D). All in all, these data suggest that LSD1 treatment of CTL *ex vivo* together with anti–PDL1 enhances long-term persistence and survival in the context of ACT, improving its clinical efficacy. To investigate whether the combined therapy LSD1i-ACT and anti-PDL1 could elicit tumor-specific immunological memory, mice cured of the primary tumor with LSD1i-ACT and anti-PDL1 treatment were rechallenged with B16-OVA, and naïve mice were challenged in parallel. While naïve mice developed large, progressively growing tumors (Figure 4D), no tumor growth occurred in the LSD1i-ACT and anti-PDL1 group 20 days after tumor rechallenge, indicating the establishment of a memory response upon tumor antigen recognition which ensures immune-surveillance.

**Figure 4.**
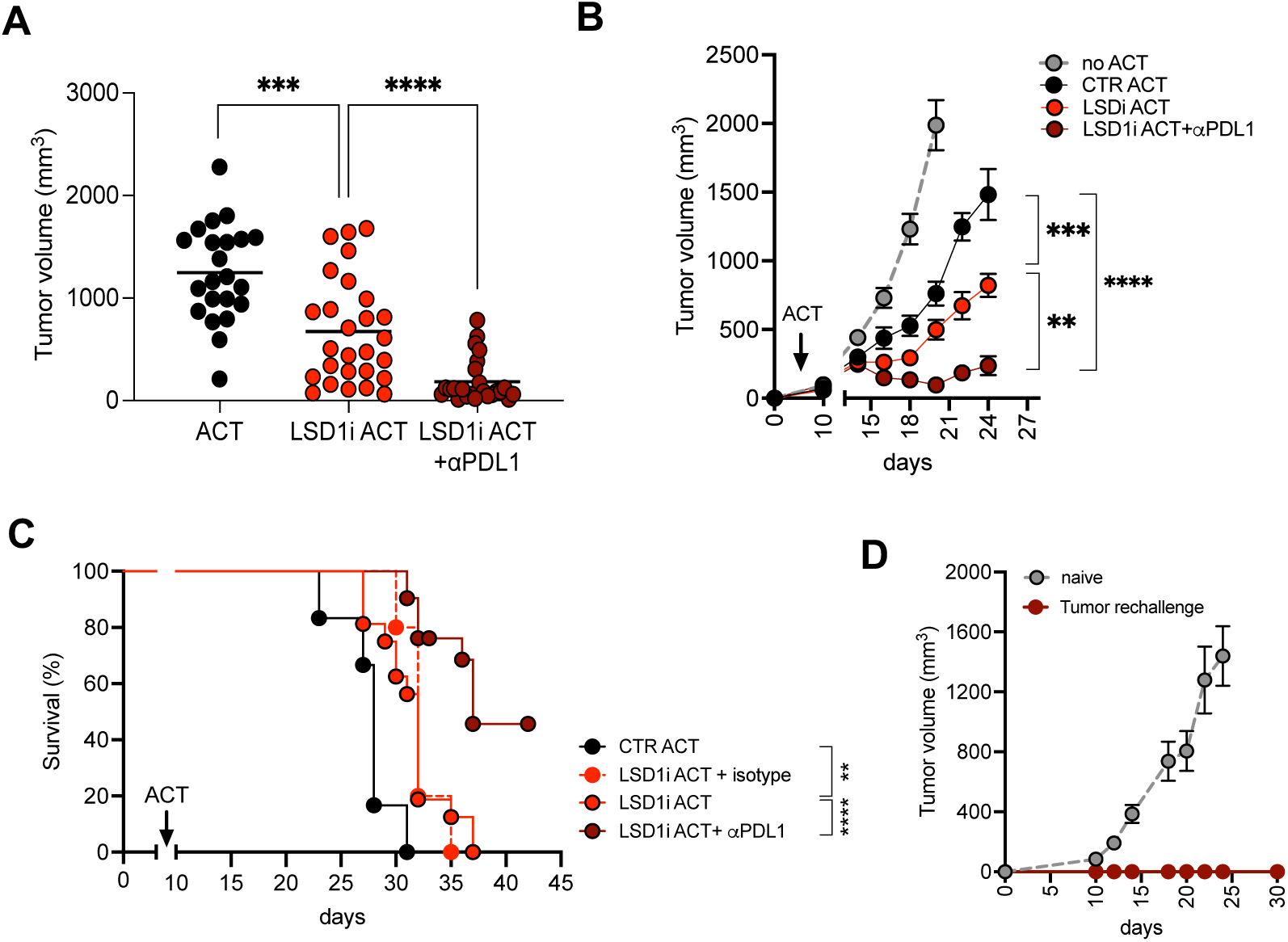
LSD1 inhibition and anti-PDL1 increases ACT efficacy. **A.** Tumor volume at the time of sacrifice, **B.** tumor growth curve and **C.** survival for CTR-ACT (n=22), LSD1i-ACT (n=27), and LSD1i-ACT+anti-PDL1 (n=27) therapy. Tumor rechallenge experiment in mice that rejected the tumor after LSD1i-ACT+anti-PDL1 therapy (D, n=8). Naïve mice were injected as controls.

All in all, this study found that, in addition to promoting the expansion, tumor infiltration and effector functions of the transferred CTL, the combined therapy LSD1i-ACT and anti-PDL1 triggers an efficient and sustainable antitumor immune response. We determined that the ACT with CTL pre-treated *in vitro* with LSD1i contributed to rapid destruction of large tumor masses while anti-PDL1 therapy concurrently prevented the emergence of a terminal exhausted phenotype that would blunts a durable antitumoral response. Indeed, only the combination of the LSD1i-ACT and anti-PDL1 secured long-term memory and disease-free survival. Our findings suggest that this combination strategy may exploit the full potential of ACT to eliminate the primary tumor, prevent immune escape, and provide long-term protective memory.

### Systemic *in vivo* LSD1i promotes intra-tumoral infiltration of memory-like CTL and increases the effectiveness of anti–PDL1 therapy

To further prove the role of LSD1i in promoting CTL anti-tumor responses and by that enhancing the efficacy of immune-based therapies, we used a preclinical mouse model of melanoma known to poorly respond to immunotherapy. BP melanoma (White et al., 2021) cell lines were implanted in mice and randomly assigned to two groups, treated or not with systemic LSD1i (DDP_38003), after establishment of the tumors (schematic representation in Figure 5A). In this experimental setting, LSD1 was inhibited systemically using the drug DDP_38003 as MC_2580 cannot be used *in vivo* (Ravasio et al., 2020). Alongside the above detected migratory signature (Figure 1B), we observed that *in vivo* systemic LSD1i promotes intra-tumoral CTL infiltration as assessed by the increased frequencies of CTL within the TME (Figure 5B). Moreover, CTL infiltrating tumors from mice treated with LSD1i display an augmented expression of Eomes (Figure 5C), which indeed translates in higher frequencies of CD44^high^/CD62L^high^ CTL (Figure 5D). Additionally, CTL from LSD1i-treated mice hold lower levels of PD1 expression (Figure 5E), further proving the role of LSD1i in pushing the polarization of CTL toward a memory-like phenotype. Nevertheless, BP tumors showed identical growth kinetic regardless LSD1i treatment (Figure 5F and S4A, C). Consistently with the observation that anti-PDL1 potentiate the anti-tumor effect of LSD1i, we decided to interrogate the role of LSD1i in combination with aPDL1 checkpoint blockade therapy to test whether the synergy between LSD1i and anti-PDL1 therapy will complement each other’s deficiencies and produce complete tumor response in tumor still refractory to immunotherapy. To this end, mice with a defined tumor mass were randomly assigned to two groups that were treated or not with anti-PDL1 therapy (Figure 5G). As expected, nor LSD1i neither anti-PDL1 treatment have any curative effect on tumor growth, while the combination between LSD1i and anti-PDL1 therapy promoted a better tumor control compared to the single treatment alone (Figure 5H and S4A-D). Interestingly, when LSD1i therapy was combined with another type of checkpoint blockade therapy - such as anti-TIM3- we did not observe any therapeutic improvements (Figure S4E), indicating a peculiar role for LSD1i in specifically improving the efficacy of anti-PDL1 based therapy.

**Figure 5.**
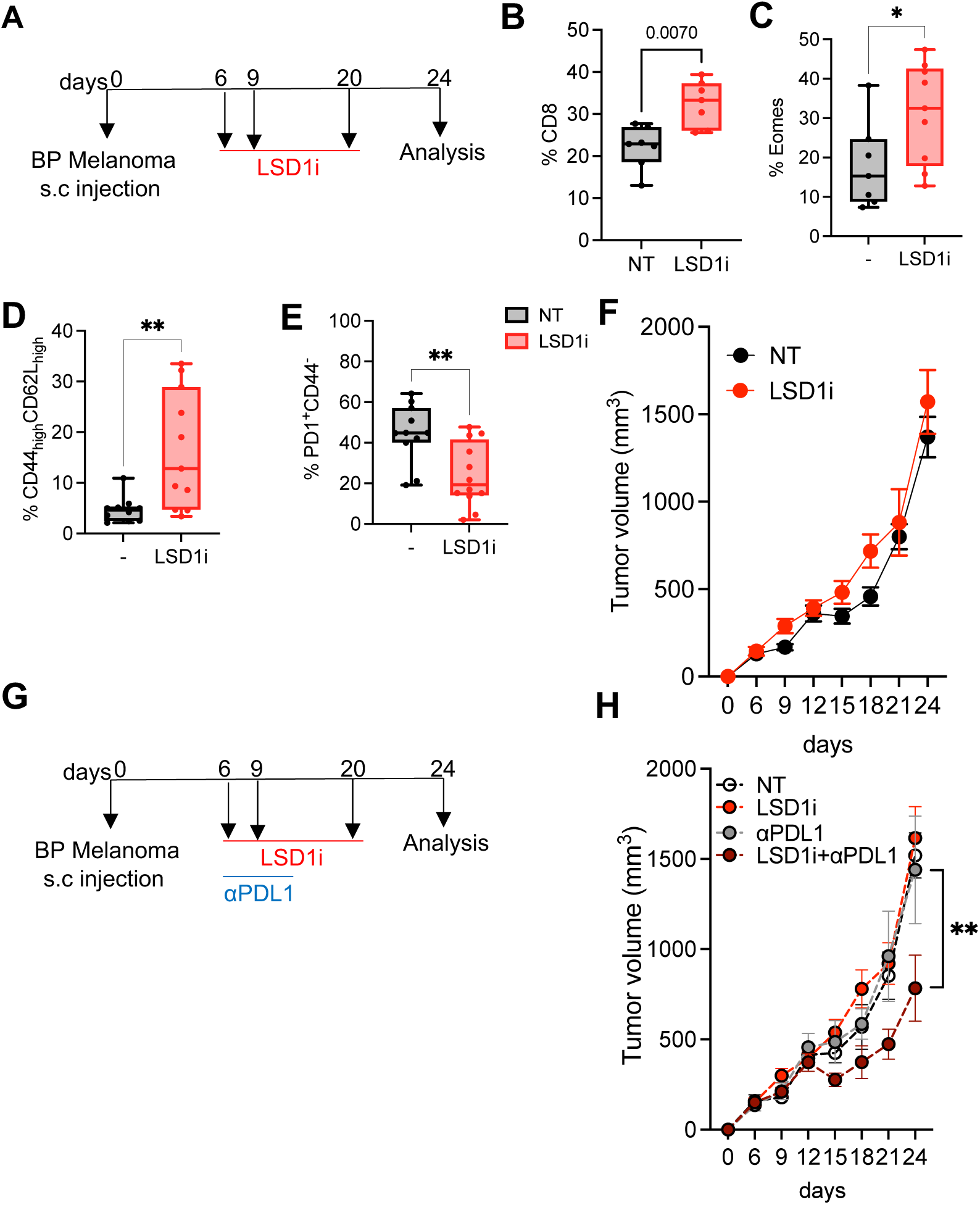
Systemic *in vivo* LSD1i promotes intra-tumoral infiltration of memory CTL and improves anti-PDL1 increases ACT efficacy. **A.** Experimental design of systemic LSD1i therapy. **B-E.** Analysis of the CTL infiltrate at the time of sacrifice. Frequencies of CTL infiltrating the tumor (B), Eomes expression (C) percentage of memory and effector cells (D) and PD1^+^/ CD44^-^ CTL (E). **F.** Tumor growth curve. **G-H.** Experimental scheme (G) and tumor growth (H, n = 11 for NT, 14 for LSD1i, 7 for anti-PDL1 and 15 for LSD1i+anti-PDL1.

In conclusion, we demonstrated the direct role of LSD1i in mediating a CTL reprogramming that improved antitumor immunity, especially in combination with anti-PDL1 therapy. LSD1i leads to an optimal priming of CTL which ameliorates their functional performances and empowers them with improved ability to debulk tumors. Its combination with anti-PDL1 therapy resulted in a complete tumor eradication and long-lasting tumor-free survival, demonstrated that LSD1i together with anti-PDL1 therapy complement each other’s deficiencies and produce a better tumor response in a melanoma model, in which both immune and epigenetic therapy alone have shown limited efficacy. Thus, this study provides a strong rationale to test a combined LSD1i- and immune-therapy in melanoma patients with primary or adaptive resistance to immunotherapy.

## METHODS

### Experimental model and subject details

Mice were housed and bred in a specific-pathogen-free animal facility and treated in accordance with the European Union Guideline on Animal Experiments under the protocol number 720/2019. For mouse experiments, we used age-matched (8-10 weeks) C57B6 or OT1 female mice in each experiment. The number of animals (biological replicates) was indicated in the respective figure legends. LCMV-P14 TCR transgenic mice were obtained through the Swiss Immunological Mouse Repository (SwImMR, Zurich, Switzerland).

### Compounds

MC_2580 and DDP_38003 have been synthesized as previously described (Vianello et al., 2016; Binda et al., 2010)

### Cell lines

BP and B16-OVA were maintained in DMEM supplemented with 10% fetal bovine serum (FBS), 1% Glutamine, 1% Pen/Strep. B16-F10 and B16OVA in complete RPMI supplemented with 10% fetal bovine serum (FBS), 1% Glutamine, 1% Pen/Strep and additional G418 (0,4mg) antibiotic for the selection and isolation of OVA-transfected B16. Cells were cultured in a humidified environment at 37 °C in 5% CO2.

### Killing experiment

For killing experiments, 2.5×10^4^ of B16-OVA cells were plated on a 96-well plate in complete RPMI media (10% FBS, 1% NaPy, 1% Glutamine, 1% Pen/Strep, 0,1% β-mercaptoethanol) supplemented with 100U/mL IL2 to adhere prior to co-culture with activated OT1 CD8+ T cells. OT1 CTL activated in the presence or absence of LSD1i for 72 hours were then counted and added to the plate in a Effector: Target (E:T) ratio of 1:1. 2,12 and 24 hours post-co-culture, cells were stained, fixed and permeabilized following manufacturer’s instruction for multiparametric flow cytometry analysis.

### CD8+ T cell isolation and activation

Spleen and lymph nodes (inguinal and axillary) were harvested from mice under sterile conditions, mechanically dissociated into single cell suspension and red blood cells were lysed. Cells were then sorted by negative selection following the manufacturer’s instructions. After sorting, cells were resuspended at 1×10^6^ cells/mL in complete RPMI media (10% FBS, 1% NaPy, 1% Glutamine, 1% Pen/Strep, 0,1% β-mercaptoethanol) supplemented with 100U/mL IL2. For OT1 and P14 CD8^+^ T cells, media was further complemented with 2 μg/mL of SIINFELK or GP-33 peptides respectively, while for C57B6 mice polyclonal activation was achieved using 5 μg/mL plated-bound αCD3 and 0,5 μg/mL soluble αCD28. Cells were further treated with LSD1 inhibitor (2 μM) for 72 hours at 37°C, 5% CO2 and 90-95% humidity for a pH neutral environment.

### Immunophenotype and flow cytometry

Characterization of the activation and differentiation profile of CTL was performed by flow cytometry following standard protocols. Briefly, for staining of surface markers 1×10^5^ – 2×10^6^ cells were harvested at the specified post-activation time points, washed with PBS, resuspended in FACS buffer (PBS, 2% FBS, EDTA 2mM) and stained for 20 minutes at 4°C. Intracellular staining for cytokine production was performed following a 4 hours incubation with complete RPMI and 1:1000 dilution of GlogiPlug in the presence or absence of a secondary stimulation with SIINFELK peptide (5 μg/mL) or PMA/Ionomycin (20 ng/mL and 1 μg/mL respectively) at 37°C. Cells were fixed and permeabilized following manufacturer’s instruction. Intracellular staining was then performed in permeabilization buffer for 2 hours at 4°C. CFSE and MitoTRK Orange were performed as per manufacturer’s instructions. Samples were acquired with BD FACS Celesta.

### tSNE analysis

FCS data was analysed with software FlowJo v10.2. RPhenograph package was used to perform computational analysis of multiparametric flow cytometry data. 3000 events per sample were concatenated by the ‘‘cytof_exprsMerge function’’, after manual gating isolation of singlet, LD negative CD45+CD3+CD8+ T cells. All samples were concatenated by the ‘‘cytof_exprsMerge function’’. /). The number of nearest neighbours identified in the first iteration of the algorithm, K value, was set to 100. UMAP and tSNE representation were generated and visualized using FlowJo version 10.2. Under-represented clusters (<0.5%) were discarded in subsequent analysis.

### Seahorse extracellular flux analysis

Seahorse experiments were performed on sorted CD8+ cells activated in presence or absence of LSDi using XF Cell Mito Stress kit (Seahorse Bioscience). OCR and ECAR were measured with XF96 Extracellular Flux Analyzers (Seahorse Bioscience). Briefly, cells were plated on poly-D-lysine–coated 96-well polystyrene Seahorse plates (200,000 T cells/well), equilibrated for 1 h at 37°C, and assayed for OCR (pmol/min) and ECAR (mpH/min) in basal conditions and after addition of oligomycin (1 μM), carbonyl cyanide-4-phenylhydrazone(1.5 μM), and antimycin A/rotenone (1 μM/0.1 μM).

### RNA sequencing and data analysis

CD8+ cells were activated for 72h in presence or absence of LSDi, counted and resuspended in RTL lysis buffer (Qiagen). RNA was isolated from purified cells using RNeasy Mini Kits, and RNA concentrations were determined using Nanodrop. Raw reads were mapped to the mouse reference genome mm9 using STAR (Dobin et al., 2013). Reads quantification was calculated using the *featureCount* function of the Subread package (Liao et al., 2014). Batch correction of the CTL-LSDi samples was assessed with the R package sva (Leek et al., 2023). *edgeR* was used to assess differential expression (Robinson et al., 2009). Differentially expressed genes (DEGs) were defined as those showing *FDR* ≤0.05 and linear fold-change ≥1.2. Pathway analysis and transcription factor prediction analysis were performed with QIAGEN’s Ingenuity Pathway Analysis (IPA, QIAGEN Redwood City, www.qiagen.com/ingenuity).

### In vivo animal experiments

C57B6 were subcutaneously injected with 2× 10^5^ B16-OVA\BP\B16-F10 cells/mouse. Animals were euthanized when the tumor reached the volume of <1 cm^3^ and >1 cm^3^ to evaluate CD8+ T cell infiltration during progression. Animal were euthanized when tumors become ulcerated or interfere with the ability of the animal to eat, drink, or ambulate. Spleens and tumors harvested and dissociated to a single cell suspension. Tumors were digested (1 mg/mL Collagenase A and 0.1 mg/mL DNase I) in DMEM at 37 °C. Cell suspension was filtered first through 100 μm and secondly through 70 μm cell strainers, washed, counted, and stained for multiparametric flow cytometry. All mouse experiments were conducted in agreement with requirements permitted by our ethical committee.

For in vivo experiments, checkpoint blockade was performed by intraperitoneal injection of 200 ug anti-mouse PD-L1 (clone 10F.9G2, Leinco Technologies) or isotype control anti-rat IgG2gb, Kappa immunoglobulin (clone R1371, Leinco Technologies) based on the following treatment schedule: five days after ACT and every other day (day 15\17\19),i.p. for three times. For indicated experiment C57B6 were rechallenged with 2 × 10^5^ B16-OVA cells/mouse 3 days after the last anti-mouse PD-L1 injection (day21).

For in vivo systemic LSD1 inhibition LSD1i DDP_38003 at the dose of 34 mg/kg was administered by oral gavage twice a week once tumors had reached a measurable size for 3 times.

### Adoptive cell transfer (ACT) experiments

*In vivo* experiments were performed using antigen specific melanoma cell line B16-OVA that was maintained in culture in RPMI, 10% FBS, 1%PenStrep, 0,4mg G418 up to 5 passages prior to injection. C57B6 mice were subcutaneously injected with 2×10^5^ cells of B16-OVA cells and tumor growth was monitored daily. 10 days post-injections and once tumors had reached a measurable size, ACT was performed intravenously using 5×10^6^ OT1 cells previously activated in the presence or absence of LSD1 inhibitor as previously described. Tumors were inspected and measured daily to construct a growth curve using the formula: Tumor volume= length x width^2^/2, where length represents the largest tumor diameter and width represents the perpendicular tumor diameter. On the day of analysis spleen and tumor were collected from each mouse, disaggregated to single cell suspension, and used for flow cytometry analysis.

### Statistical analysis and data quantification

Flow cytometry data was analyzed using FlowJo v10.2. All statistical analyses were conducted using GraphPad Prism version 9.2.0. Differences between two groups were calculated by two-tailed Student’s t test, unless otherwise specified. For graphs with multiple comparisons being made, one-way ANOVA was performed with post hoc Sidak’s test or Tukey’s test for multiple comparisons, unless otherwise specified. Significance was set at p values ≤ 0.05. For all figures: *, p ≤ 0.05; **, p ≤ 0.01; ***, p ≤ 0.001. All p values are indicated in the figure legend. All plots indicate Min to Max value data and error bars are shown as mean ± SEM. The n numbers for each experiment and the numbers of experiments are annotated in each figure legends. For all experiments, data are reported based on individual biological replicates pooled from multiple donors or animals.

## ACKNOWLEDGMENTS

The authors declare that they have no competing interests.

We thank IEO’s Genomic Unit, Flow Cytometry and Imaging Unit at IEO for their technical assistance. This work was supported by Associazione Italiana per la Ricerca contro il Cancro (AIRC StartUp Grant 21474 to TM, AIRC IG 20 to SM). This work was partially supported by the Italian Ministry of Health with Ricerca Corrente and 5 x 1000 funds. TM thanks Leonardo and Lara Nezi for their support.

---

**Figure Supplementary 1 (related to Figure 1).**
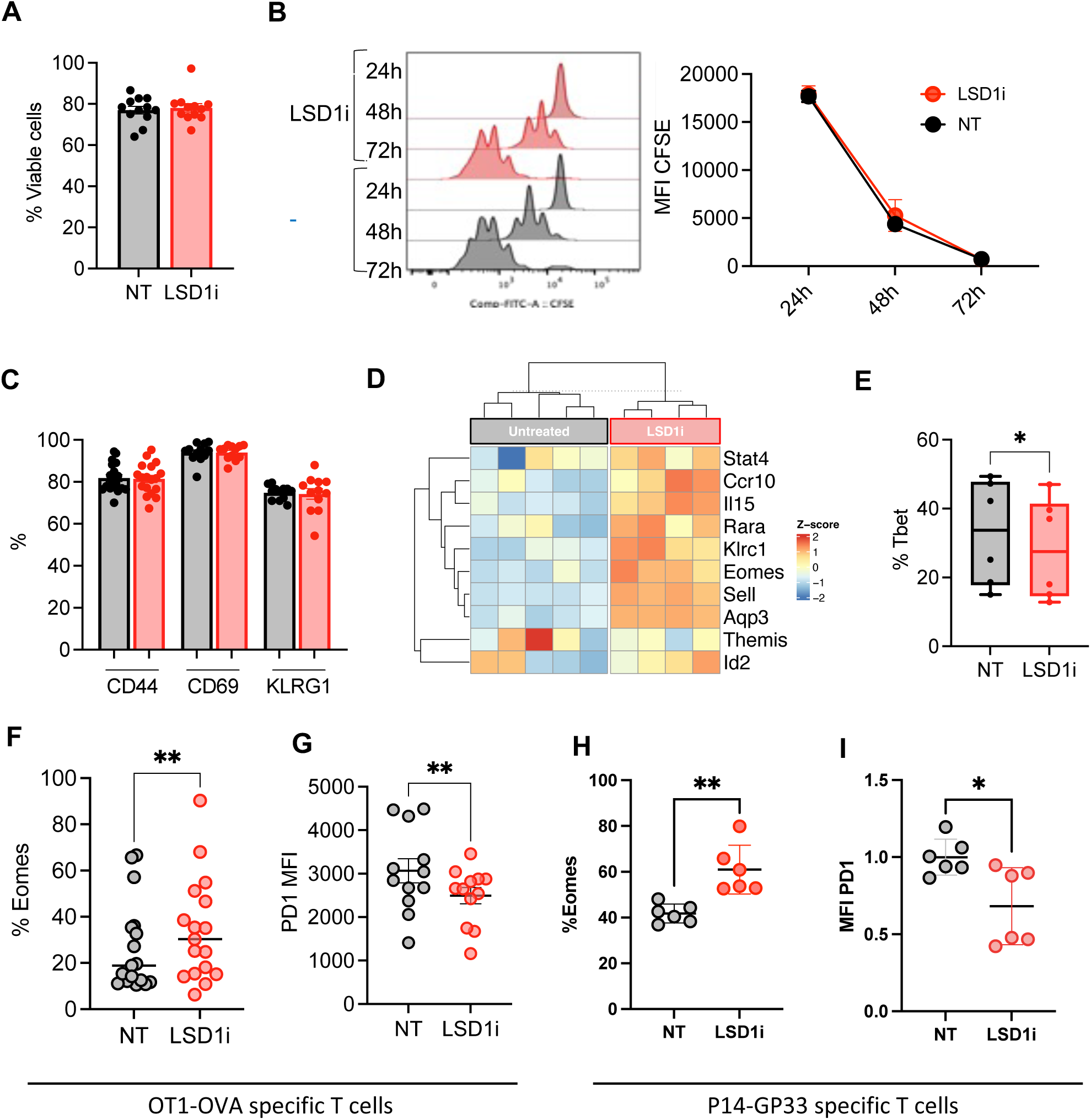
**A.** Percentage of viable cells in CTL±LSD1i (n=12)**. B.** Proliferation ability measured by CFSE profile at different time points (24,48,72h) **C.** Activation status measures as canonical activation markers (CD44,CD69,KLRG1, n=12). **D.** Heatmap showing expression of selected memory-related genes in CTL±LSD1i. **E.** Tbet expression levels (n=8). **F-I.** LSD1i effect on antigen-primed CTL from OT1 (F-G, n=12) and P14 (H-I, n=6) mice. Data from 4/16 biological replicates; 2–5 independent experiments.

**Figure Supplementary 2 (related to Figure 2).**
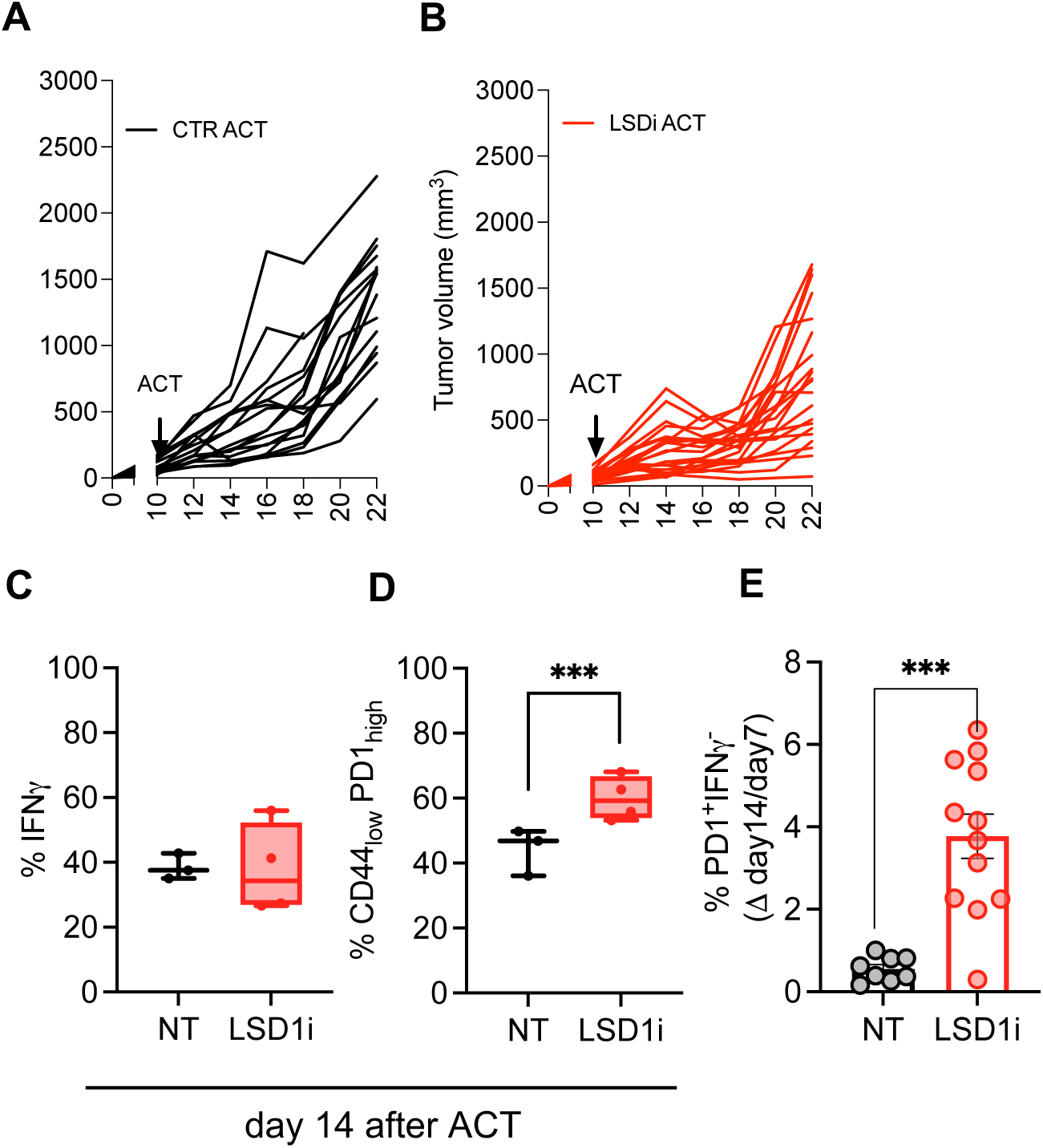
**A-B.** Spider plots representing tumor growth curves for CTR-ACT (A) and LSD1i-ACT (B). **C-E.** Analysis of the CTL infiltrate 14 days post-infusion. Frequencies of IFNγ−secreting CTL (C) and exhaustion profile as PD1_high_CD44_low_ (D) and PD1_high_IFN_low_ CTL (E).

**Figure Supplementary 3 (related to Figure 4).**
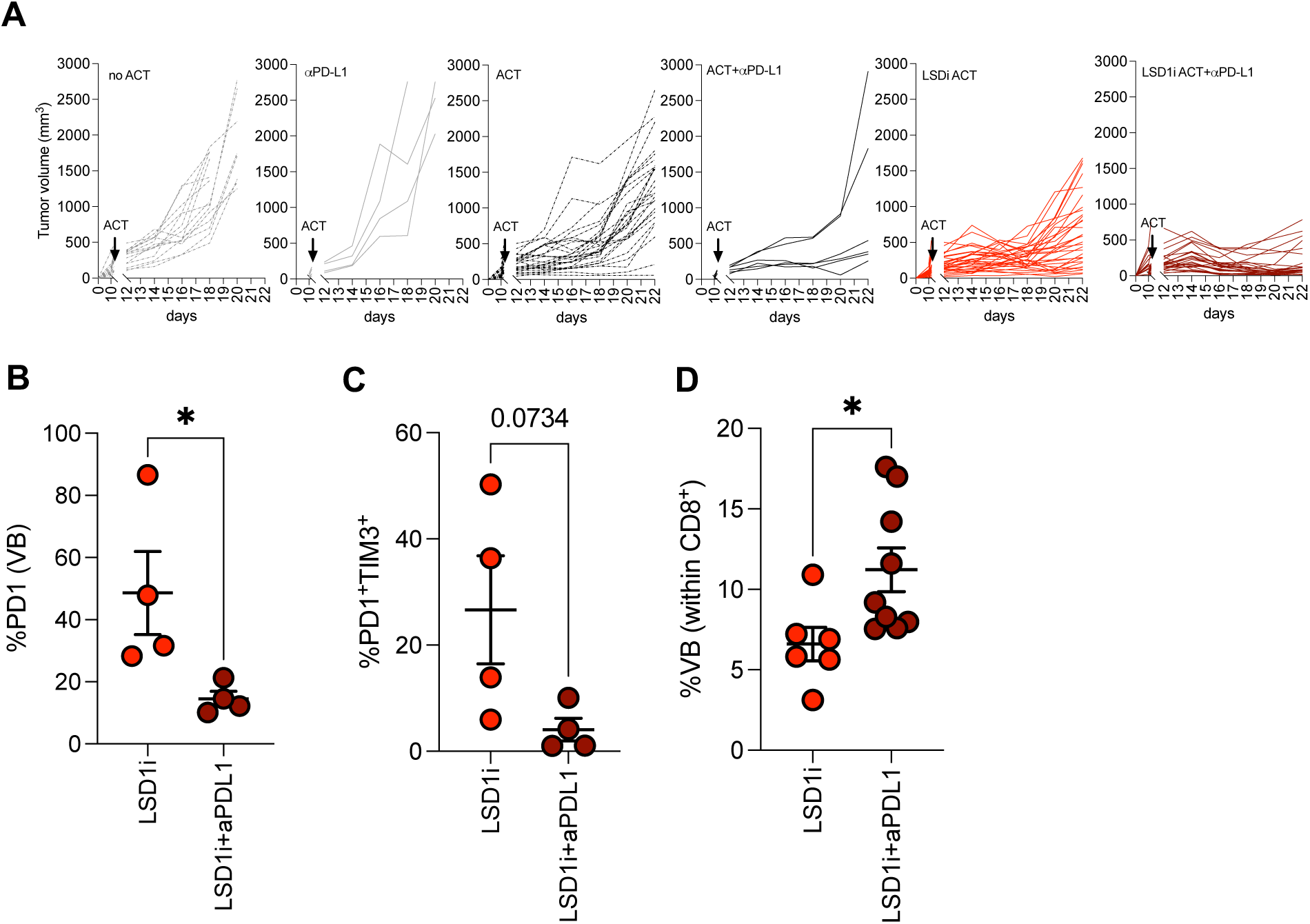
**A.** Spider plots representing tumor growth curves for the indicated treatments. **B-D.** Analysis of the CTL infiltrate at the time of sacrifice. Frequencies of PD1_pos_ (B, n=4) and exhaustion profile as PD1_pos_TIM3_pos_ (C, n=4) and Vbeta5.2 CTL (D, n=6) .

**Figure Supplementary 4 (related to Figure 5).**
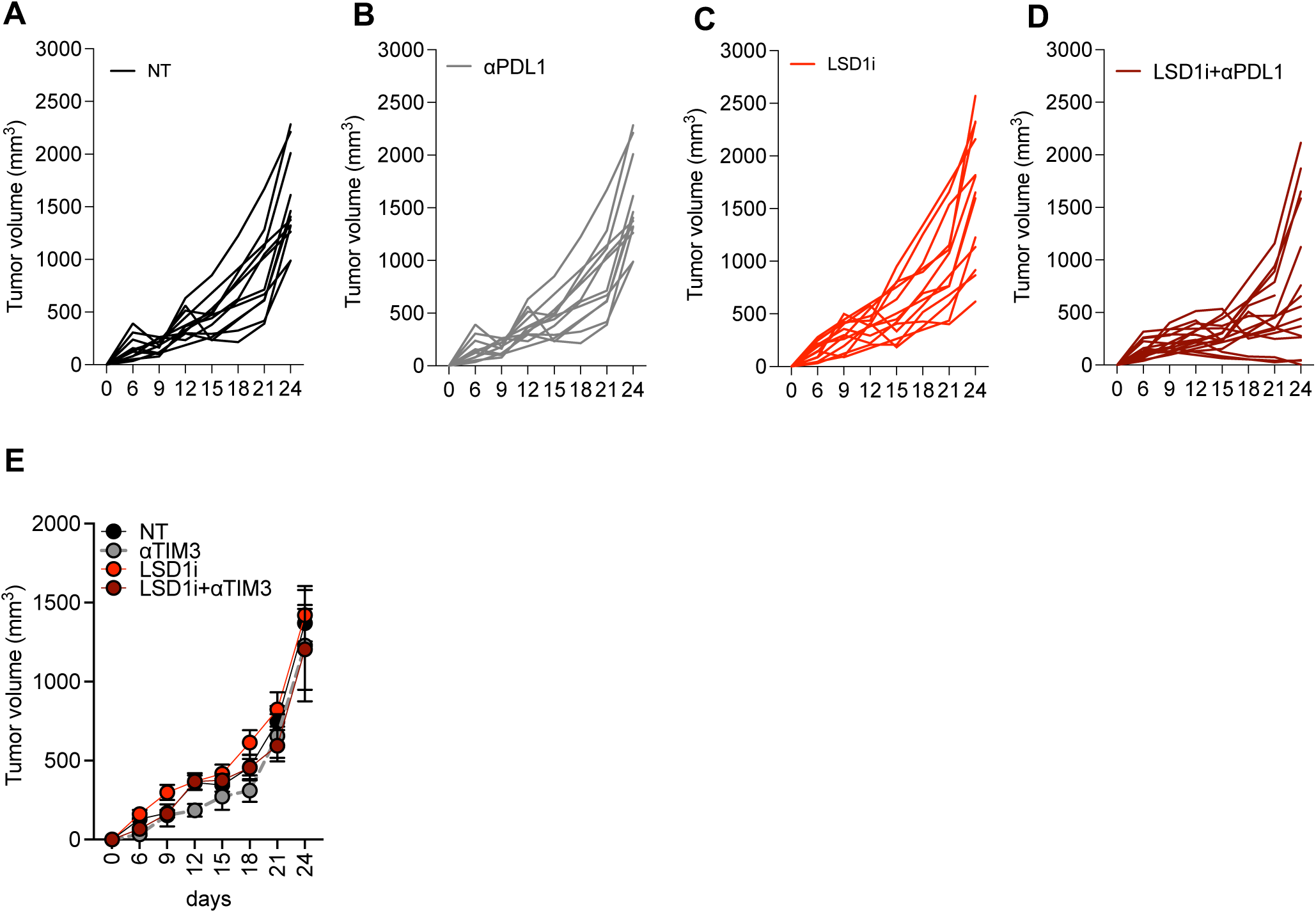
**A-D.** Spider plots representing tumor growth curves for the indicated treatments. **E.** Analysis of the tumor growth curves for LSD1i and anti-TIM3 alone or in combination.

## REFERENCES

Benci, J.L., B. Xu, Y. Qiu, T.J. Wu, H. Dada, C. Twyman-Saint Victor, L. Cucolo, D.S.M. Lee, K.E. Pauken, A.C. Huang, T.C. Gangadhar, R.K. Amaravadi, L.M. Schuchter, M.D. Feldman, H. Ishwaran, R.H. Vonderheide, A. Maity, E.J. Wherry, and A.J. Minn. 2016. Tumor Interferon Signaling Regulates a Multigenic Resistance Program to Immune Checkpoint Blockade. Cell. 167:1540–1554.e12. doi:10.1016/j.cell.2016.11.022.

Binda, C., S. Valente, M. Romanenghi, S. Pilotto, R. Cirilli, A. Karytinos, G. Ciossani, O.A. Botrugno, F. Forneris, M. Tardugno, D.E. Edmondson, S. Minucci, A. Mattevi, and A. Mai. 2010. Biochemical, structural, and biological evaluation of tranylcypromine derivatives as inhibitors of histone demethylases LSD1 and LSD2. J Am Chem Soc. 132:6827–6833. doi:10.1021/ja101557k.

Brentjens, R.J., M.L. Davila, I. Riviere, J. Park, X. Wang, L.G. Cowell, S. Bartido, J. Stefanski, C. Taylor, M. Olszewska, O. Borquez-Ojeda, J. Qu, T. Wasielewska, Q. He, Y. Bernal, I. V. Rijo, C. Hedvat, R. Kobos, K. Curran, P. Steinherz, J. Jurcic, T. Rosenblat, P. Maslak, M. Frattini, and M. Sadelain. 2013. CD19-targeted T cells rapidly induce molecular remissions in adults with chemotherapy-refractory acute lymphoblastic leukemia. Sci Transl Med. 5. doi:10.1126/scitranslmed.3005930.

Dobin, A., C.A. Davis, F. Schlesinger, J. Drenkow, C. Zaleski, S. Jha, P. Batut, M. Chaisson, and T.R. Gingeras. 2013. STAR: Ultrafast universal RNA-seq aligner. Bioinformatics. 29:15–21. doi:10.1093/bioinformatics/bts635.

Doering, T.A., A. Crawford, J.M. Angelosanto, M.A. Paley, C.G. Ziegler, and E.J. Wherry. 2012. Network Analysis Reveals Centrally Connected Genes and Pathways Involved in CD8+ T Cell Exhaustion versus Memory. Immunity. 37:1130–1144. doi:10.1016/j.immuni.2012.08.021.

Fraietta, J.A., C.L. Nobles, M.A. Sammons, S. Lundh, S.A. Carty, T.J. Reich, A.P. Cogdill, J.J.D. Morrissette, J.E. DeNizio, S. Reddy, Y. Hwang, M. Gohil, I. Kulikovskaya, F. Nazimuddin, M. Gupta, F. Chen, J.K. Everett, K.A. Alexander, E. Lin-Shiao, M.H. Gee, X. Liu, R.M. Young, D. Ambrose, Y. Wang, J. Xu, M.S. Jordan, K.T. Marcucci, B.L. Levine, K.C. Garcia, Y. Zhao, M. Kalos, D.L. Porter, R.M. Kohli, S.F. Lacey, S.L. Berger, F.D. Bushman, C.H. June, and J.J. Melenhorst. 2018. Disruption of TET2 promotes the therapeutic efficacy of CD19-targeted T cells. Nature. 558:307–312. doi:10.1038/s41586-018-0178-z.

Ghoneim, H.E., Y. Fan, A. Moustaki, H.A. Abdelsamed, P. Dash, P. Dogra, R. Carter, W. Awad, G. Neale, P.G. Thomas, and B. Youngblood. 2017. De Novo Epigenetic Programs Inhibit PD-1 Blockade-Mediated T Cell Rejuvenation. Cell. 170:142–157.e19. doi:10.1016/j.cell.2017.06.007.

Goswami, S., I. Apostolou, J. Zhang, J. Skepner, S. Anandhan, X. Zhang, L. Xiong, P. Trojer, A. Aparicio, S.K. Subudhi, J.P. Allison, H. Zhao, and P. Sharma. 2018. Modulation of EZH2 expression in T cells improves efficacy of anti-CTLA-4 therapy. Journal of Clinical Investigation. 128:3813–3818. doi:10.1172/JCI99760.

Grasso, C.S., J. Tsoi, M. Onyshchenko, G. Abril-Rodriguez, P. Ross-Macdonald, M. Wind-Rotolo, A. Champhekar, E. Medina, D.Y. Torrejon, D.S. Shin, P. Tran, Y.J. Kim, C. Puig-Saus, K. Campbell, A. Vega-Crespo, M. Quist, C. Martignier, J.J. Luke, J.D. Wolchok, D.B. Johnson, B. Chmielowski, F.S. Hodi, S. Bhatia, W. Sharfman, W.J. Urba, C.L. Slingluff, A. Diab, J.B.A.G. Haanen, S.M. Algarra, D.M. Pardoll, V. Anagnostou, S.L. Topalian, V.E. Velculescu, D.E. Speiser, A. Kalbasi, and A. Ribas. 2020. Conserved Interferon-γ Signaling Drives Clinical Response to Immune Checkpoint Blockade Therapy in Melanoma. Cancer Cell. 38:500–515.e3. doi:10.1016/j.ccell.2020.08.005.

Gray, S.M., R.A. Amezquita, T. Guan, S.H. Kleinstein, and S.M. Kaech. 2017. Polycomb Repressive Complex 2-Mediated Chromatin Repression Guides Effector CD8+ T Cell Terminal Differentiation and Loss of Multipotency. Immunity. 46:596–608. doi:10.1016/j.immuni.2017.03.012.

Hoekstra, M.E., L. Bornes, F.E. Dijkgraaf, D. Philips, I.N. Pardieck, M. Toebes, D.S. Thommen, J. van Rheenen, and T.N.M. Schumacher. 2020. Long-distance modulation of bystander tumor cells by CD8+ T-cell-secreted IFN-γ. Nat Cancer. 1:291–301. doi:10.1038/s43018-020-0036-4.

Hosseini, A., and S. Minucci. 2017. A comprehensive review of lysine-specific demethylase 1 and its roles in cancer. Epigenomics. 9:1123–1142. doi:10.2217/epi-2017-0022.

Kalos, M., B.L. Levine, D.L. Porter, S. Katz, S.A. Grupp, A. Bagg, and C.H. June. 2011. T cells with chimeric antigen receptors have potent antitumor effects and can establish memory in patients with advanced leukemia. Sci Transl Med. 3:1–12. doi:10.1126/scitranslmed.3002842.

Kishton, R.J., M. Sukumar, and N.P. Restifo. 2017. Metabolic Regulation of T Cell Longevity and Function in Tumor Immunotherapy. Cell Metab. 26:94–109. doi:10.1016/j.cmet.2017.06.016.

Klebanoff, C.A., L. Gattinoni, and N.P. Restifo. 2012. Sorting through subsets: Which T-cell populations mediate highly effective adoptive immunotherapy? Journal of Immunotherapy. 35:651–660. doi:10.1097/CJI.0b013e31827806e6.

Kuwahara, M., M. Yamashita, K. Shinoda, S. Tofukuji, A. Onodera, R. Shinnakasu, S. Motohashi, H. Hosokawa, D. Tumes, C. Iwamura, V. Lefebvre, and T. Nakayama. 2012. The transcription factor Sox4 is a downstream target of signaling by the cytokine TGF-β and suppresses T H2 differentiation. Nat Immunol. 13:778–786. doi:10.1038/ni.2362.

Leek, A.J.T., W.E. Johnson, H.S. Parker, E.J. Fertig, A.E. Jaffe, Y. Zhang, J.D. Storey, L.C. Torres, and W.E. Johnson. 2023. Package ‘ sva .’

Liao, Y., G.K. Smyth, and W. Shi. 2014. FeatureCounts: An efficient general purpose program for assigning sequence reads to genomic features. Bioinformatics. 30:923–930. doi:10.1093/bioinformatics/btt656.

Liu, Y., B. Debo, M. Li, Z. Shi, W. Sheng, and Y. Shi. 2021. LSD1 inhibition sustains T cell invigoration with a durable response to PD-1 blockade. Nat Commun. 12:1–16. doi:10.1038/s41467-021-27179-7.

Manzo, T., B.M. Prentice, K.G. Anderson, A. Raman, A. Schalck, G.S. Codreanu, C.B. Nava Lauson, S. Tiberti, A. Raimondi, M.A. Jones, M. Reyzer, B.M. Bates, J.M. Spraggins, N.H. Patterson, J.A. McLean, K. Rai, C. Tacchetti, S. Tucci, J.A. Wargo, S. Rodighiero, K. Clise-Dwyer, S.D. Sherrod, M. Kim, N.E. Navin, R.M. Caprioli, P.D. Greenberg, G. Draetta, and L. Nezi. 2020. Accumulation of long-chain fatty acids in the tumor microenvironment drives dysfunction in intrapancreatic cd8+ t cells. Journal of Experimental Medicine. 217. doi:10.1084/jem.20191920.

Martin, M.D., and V.P. Badovinac. 2018. Defining memory CD8 T cell. Front Immunol. 9:1–10. doi:10.3389/fimmu.2018.02692.

Maude, S.L., N. Frey, P.A. Shaw, R. Aplenc, D.M. Barrett, N.J. Bunin, A. Chew, V.E. Gonzalez, Z. Zheng, S.F. Lacey, Y.D. Mahnke, J.J. Melenhorst, S.R. Rheingold, A. Shen, D.T. Teachey, B.L. Levine, C.H. June, D.L. Porter, and S.A. Grupp. 2014. Chimeric Antigen Receptor T Cells for Sustained Remissions in Leukemia. New England Journal of Medicine. 371:1507–1517. doi:10.1056/nejmoa1407222.

McLane, L.M., M.S. Abdel-Hakeem, and E.J. Wherry. 2019. CD8 T Cell Exhaustion During Chronic Viral Infection and Cancer. Annu Rev Immunol. 37:457–495. doi:10.1146/annurev-immunol-041015-055318.

Mondino, A., and T. Manzo. 2020. To Remember or to Forget: The Role of Good and Bad Memories in Adoptive T Cell Therapy for Tumors. Front Immunol. 11:1–15. doi:10.3389/fimmu.2020.01915.

Nava Lauson, C.B., S. Tiberti, P.A. Corsetto, F. Conte, P. Tyagi, M. Machwirth, S. Ebert, A. Loffreda, L. Scheller, D. Sheta, Z. Mokhtari, T. Peters, A.T. Raman, F. Greco, A.M. Rizzo, A. Beilhack, G. Signore, N. Tumino, P. Vacca, L.A. McDonnell, A. Raimondi, P.D. Greenberg, J.B. Huppa, S. Cardaci, I. Caruana, S. Rodighiero, L. Nezi, and T. Manzo. 2023. Linoleic acid potentiates CD8+ T cell metabolic fitness and antitumor immunity. Cell Metab. 35:633–650.e9. doi:10.1016/j.cmet.2023.02.013.

Paschen, A., I. Melero, and A. Ribas. 2022. Central Role of the Antigen-Presentation and Interferon-γ Pathways in Resistance to Immune Checkpoint Blockade. Annu Rev Cancer Biol. 6:85–102. doi:10.1146/annurev-cancerbio-070220-111016.

Pauken, K.E., M.A. Sammons, P.M. Odorizzi, S. Manne, J. Godec, O. Khan, A.M. Drake, Z. Chen, D.R. Sen, M. Kurachi, R.A. Barnitz, C. Bartman, B. Bengsch, A.C. Huang, J.M. Schenkel, G. Vahedi, W.N. Haining, S.L. Berger, and E.J. Wherry. 2016. Epigenetic stability of exhausted T cells limits durability of reinvigoration by PD-1 blockade. Science (1979). 354:1160–1165. doi:10.1126/science.aaf2807.

Philip, M., L. Fairchild, L. Sun, E.L. Horste, S. Camara, M. Shakiba, A.C. Scott, A. Viale, P. Lauer, T. Merghoub, M.D. Hellmann, J.D. Wolchok, C.S. Leslie, and A. Schietinger. 2017. Chromatin states define tumour-specific T cell dysfunction and reprogramming. Nature. 545:452–456. doi:10.1038/nature22367.

Qin, Y., S.N. Vasilatos, L. Chen, H. Wu, Z. Cao, Y. Fu, M. Huang, A.M. Vlad, B. Lu, S. Oesterreich, N.E. Davidson, and Y. Huang. 2019. Inhibition of histone lysine-specific demethylase 1 elicits breast tumor immunity and enhances antitumor efficacy of immune checkpoint blockade. Oncogene. 38:390–405. doi:10.1038/s41388-018-0451-5.

Rapoport, A.P., E.A. Stadtmauer, G.K. Binder-Scholl, O. Goloubeva, D.T. Vogl, S.F. Lacey, A.Z. Badros, A. Garfall, B. Weiss, J. Finklestein, I. Kulikovskaya, S.K. Sinha, S. Kronsberg, M. Gupta, S. Bond, L. Melchiori, J.E. Brewer, A.D. Bennett, A.B. Gerry, N.J. Pumphrey, D. Williams, H.K. Tayton-Martin, L. Ribeiro, T. Holdich, S. Yanovich, N. Hardy, J. Yared, N. Kerr, S. Philip, S. Westphal, D.L. Siegel, B.L. Levine, B.K. Jakobsen, M. Kalos, and C.H. June. 2015. NY-ESO-1-specific TCR-engineered T cells mediate sustained antigen-specific antitumor effects in myeloma. Nat Med. 21:914–921. doi:10.1038/nm.3910.

Ravasio, R., E. Ceccacci, L. Nicosia, A. Hosseini, P.L. Rossi, I. Barozzi, L. Fornasari, R.D. Zuffo, S. Valente, R. Fioravanti, C. Mercurio, M. Varasi, A. Mattevi, A. Mai, G. Pavesi, T. Bonaldi, and S. Minucci. 2020. Targeting the scaffolding role of LSD1 (KDM1A) poises acute myeloid leukemia cells for retinoic acid-induced differentiation. Sci Adv. 6:1–14. doi:10.1126/sciadv.aax2746.

Robinson, M.D., D.J. McCarthy, and G.K. Smyth. 2009. edgeR: A Bioconductor package for differential expression analysis of digital gene expression data. Bioinformatics. 26:139–140. doi:10.1093/bioinformatics/btp616.

Salas-Benito, D., T.R. Berger, and M. V. Maus. 2023. Stalled CARs: Mechanisms of Resistance to CAR T Cell Therapies. Annu Rev Cancer Biol. 7:23–42. doi:10.1146/annurev-cancerbio-061421-012235.

Sen, D.R., J. Kaminski, R.A. Barnitz, M. Kurachi, U. Gerdemann, K.B. Yates, H.W. Tsao, J. Godec, M.W. LaFleur, F.D. Brown, P. Tonnerre, R.T. Chung, D.C. Tully, T.M. Allen, N. Frahm, G.M. Lauer, E.J. Wherry, N. Yosef, and W.N. Haining. 2016. The epigenetic landscape of T cell exhaustion. Science (1979). 354:1165–1169. doi:10.1126/science.aae0491.

Sheng, W., M.W. LaFleur, T.H. Nguyen, S. Chen, A. Chakravarthy, J.R. Conway, Y. Li, H. Chen, H. Yang, P.H. Hsu, E.M. Van Allen, G.J. Freeman, D.D. De Carvalho, H.H. He, A.H. Sharpe, and Y. Shi. 2018. LSD1 Ablation Stimulates Anti-tumor Immunity and Enables Checkpoint Blockade. Cell. 174:549–563.e19. doi:10.1016/j.cell.2018.05.052.

Sukumar, M., R.J. Kishton, and N.P. Restifo. 2017. Metabolic reprograming of anti-tumor immunity. Curr Opin Immunol. 46:14–22. doi:10.1016/j.coi.2017.03.011.

Thibaut, R., P. Bost, I. Milo, M. Cazaux, F. Lemaître, Z. Garcia, I. Amit, B. Breart, C. Cornuot, B. Schwikowski, and P. Bousso. 2020. Bystander IFN-γ activity promotes widespread and sustained cytokine signaling altering the tumor microenvironment. Nat Cancer. 1:302–314. doi:10.1038/s43018-020-0038-2.

Trinchieri, G. 2003. Interleukin-12 and the regulation of innate resistance and adaptive immunity. Nat Rev Immunol. 3:133–146. doi:10.1038/nri1001.

Tu, W.J., R.D. McCuaig, A.H.Y. Tan, K. Hardy, N. Seddiki, S. Ali, J.E. Dahlstrom, E.G. Bean, J. Dunn, J. Forwood, S. Tsimbalyuk, K. Smith, D. Yip, L. Malik, T. Prasanna, P. Milburn, and S. Rao. 2020. Targeting Nuclear LSD1 to Reprogram Cancer Cells and Reinvigorate Exhausted T Cells via a Novel LSD1-EOMES Switch. Front Immunol. 11:1–23. doi:10.3389/fimmu.2020.01228.

Vianello, P., O.A. Botrugno, A. Cappa, R. Dal Zuffo, P. Dessanti, A. Mai, B. Marrocco, A. Mattevi, G. Meroni, S. Minucci, G. Stazi, F. Thaler, P. Trifiró, S. Valente, M. Villa, M. Varasi, and C. Mercurio. 2016. Discovery of a Novel Inhibitor of Histone Lysine-Specific Demethylase 1A (KDM1A/LSD1) as Orally Active Antitumor Agent. J Med Chem. 59:1501–1517. doi:10.1021/acs.jmedchem.5b01209.

White, M.G., R. Szczepaniak Sloane, R.G. Witt, A. Reuben, P.O. Gaudreau, M.C. Andrews, N. Feng, S. Johnson, C.A. Class, C. Bristow, K. Wani, C. Hudgens, L. Nezi, T. Manzo, M.P. De Macedo, J. Hu, R. Davis, H. Jiang, P. Prieto, E. Burton, P. Hwu, H. Tawbi, J. Gershenwald, A.J. Lazar, M.T. Tetzlaff, W. Overwijk, S.E. Woodman, Z.A. Cooper, J.R. Marszalek, M.A. Davies, T.P. Heffernan, and J.A. Wargo. 2021. Short-term treatment with multi-drug regimens combining BRAF/MEK-targeted therapy and immunotherapy results in durable responses in Braf-mutated melanoma. Oncoimmunology. 10. doi:10.1080/2162402X.2021.1992880.

van der Windt, G.J.W., B. Everts, C.H. Chang, J.D. Curtis, T.C. Freitas, E. Amiel, E.J. Pearce, and E.L. Pearce. 2012. Mitochondrial Respiratory Capacity Is a Critical Regulator of CD8 + T Cell Memory Development. Immunity. 36:68–78. doi:10.1016/j.immuni.2011.12.007.

Young, R.M., N.W. Engel, U. Uslu, N. Wellhausen, and C.H. June. 2022. Next-Generation CAR T-cell Therapies. Cancer Discov. 12:1625–1633. doi:10.1158/2159-8290.CD-21-1683.

Zaid, A., J.L. Hor, S.N. Christo, J.R. Groom, W.R. Heath, L.K. Mackay, and S.N. Mueller. 2017. Chemokine Receptor–Dependent Control of Skin Tissue–Resident Memory T Cell Formation. The Journal of Immunology. 199:2451–2459. doi:10.4049/jimmunol.1700571.

